# Somrit: The Somatic Retrotransposon Insertion Toolkit

**DOI:** 10.1101/2023.08.06.552193

**Authors:** Alister V. D’Costa, Jared T. Simpson

**Affiliations:** Ontario Institute for Cancer Research, Toronto, Canada; Department of Computer Science, University of Toronto, Toronto, Canada; Department of Molecular Genetics, University of Toronto, Toronto, Canada

**Keywords:** retrotransposon, somatic, structural variation, nanopore

## Abstract

Mobile elements, such as retrotransposons, have the ability to express and re-insert themselves into the genome, with over half the human genome being made up of mobile element sequence. Somatic mobile element insertions (MEIs) have been shown to cause disease, including some cancers. Accurate identification of where novel retrotransposon insertion events occur in the genome is crucial to understand the functional consequence of an insertion event. In this paper we describe somrit, a modular toolkit for detecting somatic MEIs from long reads aligned to a reference genome. We identify the initial read-to-reference mapping step as a potential source of error when the insertion is similar to a nearby repeat in the reference genome and develop a consensus-realignment procedure to resolve this. We show how somrit improves the sensitivity of detection for rare somatic retrotransposon insertion events compared to existing tools, and how the local realignment procedure can reduce false positive translocation calls caused by mis-mapped reads bearing MEIs. Somrit is openly available at: https://github.com/adcosta17/somrit

## Background

Mobile elements are DNA sequences that can change genomic position and re-insert themselves into the genome [1]. A large fraction of the human genome is composed of mobile element sequence with thousands of identified copies [1, 2, 3]. While many of these copies are partial fragments reflecting ancient insertion events that can no longer actively move [2, 3, 4], some more recent copies retain the ability to be expressed and re-insert themselves [5, 6]. Retrotransposons are a class of mobile elements that includes LINE-1 (L1), Alu and SVA elements [7, 8, 9]. Full length human LINE-1 elements are *∼*6kbp in length and encode proteins for retrotrans-position, allowing for their re-insertion into the genome via an RNA intermediate and reverse transcription [10, 11]. Smaller SVA (*∼*2000bp) and Alu (*∼*300bp) elements rely on the LINE-1 retrotransposition mechanism for re-integration into the genome [12, 13, 14]. LINE-1 insertions usually occur at LINE-1 endonuclease recognition motifs [15, 16], often include a target-site duplication (TSD) [15, 17] and a poly-A tail [18] and may contain genomic flaking sequence from the LINE-1 element of origin [19, 20]. Due to their mobile nature the exact number and location of retrotransposons in the genome varies from person to person, with any individual having some inherited copies not found in the human reference, and possibly somatic copies present in a subset of cells [21].

While often only a handful of retrotransposon copies in any given individual retain the ability to be expressed and re-inserted back into the genome, their expression has been linked to disease progression. Prior research has shown somatic insertion of LINE-1 elements activates oncogenes and directly drives cancer progression in some colorectal cancers [22]. Somatic insertion of LINE-1 elements may alter gene expression, including a slowing of DNA translation possibly affecting the expression of tumor suppressor genes [10]. Due to the large amount of mobile element sequence in the genome, retrotransposon insertions have the potential to generate chromosomal rearrangements including deletions, duplications, inversions and translocations, as they may mislead homologous recombination repair pathways to cause non-allelic homologous recombination events [23, 24]. These larger changes in the genome can contribute to the loss of tumor suppressors, activation of oncogenes or the generation of fusion proteins that may drive cancer progression [25].

Tools for detecting germline and somatic mobile element insertions (MEIs), including retrotransposons, have been developed for short reads including MELT[26], TraFiC-mem [27], RetroSeq [28] and xTea [29]. While effective, short reads have limited repeat resolution for large insertions, insertions containing varying numbers of repeat copies and insertions into existing repetitive sequence being particularly problematic and hard to detect [30, 31, 32]. Long read technologies like the Oxford Nanopore (ONT) and Pacific Biosciences (PacBio) instruments can generate sequencing reads exceeding 10kbp. These reads can therefore fully span a retrotransposon insertion with flanking sequence allowing the genomic location of the insertion to be identified [33] (e.g. a full length *∼*6kbp LINE-1 element can be fully contained within a 10kbp read with 4kb of flanking sequence available to inform the location of the repeat). This has prompted the development of tools to detect mobile element insertions from long reads such as tldr [34] and xTea-Long [29]. These tools have mainly been designed to detect polymorphic repeats that present as heterozygous and homozygous variants within an individual genome and hence they often require multiple reads to support an insertion call. When looking at somatic variation, such as in a tumor, insertions may occur at very low frequencies and hence be supported by only a single (or very few) reads depending on the variant allele frequency within the cellular population. Methods designed to detect polymorphic variation may miss these somatic insertion events due to their very low read support. Additionally, many large insertion events in long reads may not be correctly identified by current state of the art long read aligners.

While *de novo* assembly of diploid genomes is becoming the gold-standard method for detecting structural variants, including retrotransposon insertions, most methods currently rely on mapping reads to a reference genome [35]. Hence, having high quality read-reference alignments is crucial to detecting MEIs. Aligners such as minimap2 [36] are designed to tolerate large gaps in the alignment by using affine gap-scoring penalties [36, 37, 38, 39]. Despite these scoring schemes, the alignments may be truncated just prior to the insertion event, lowering the read support for the insertion. Further, we have found that the location of the insertion event introduced by the aligner may differ read-to-read when the inserted sequence is similar to a repeat copy already existing in the reference, an effect we term *repeat alignment ambiguity*, and also observed by [40]. This problem is analogous to the classical case of aligning two sequences with different lengths of homopolymer runs. In that case, the exact base that has been inserted/deleted is not known, so the placement of the gap is ambiguous with multiple alignments having the same alignment score (illustrated in **Figure 1B**). In our case, the repeat could be placed either before or after the existing element and the aligner’s choice may depend solely on the pattern of matches/mismatches caused by sequencing errors (**Figure 1C**). Later in the Results section, we quantify how often this artifact occurs as a function sequence divergence between the repetitive elements.

**Figure 1.**
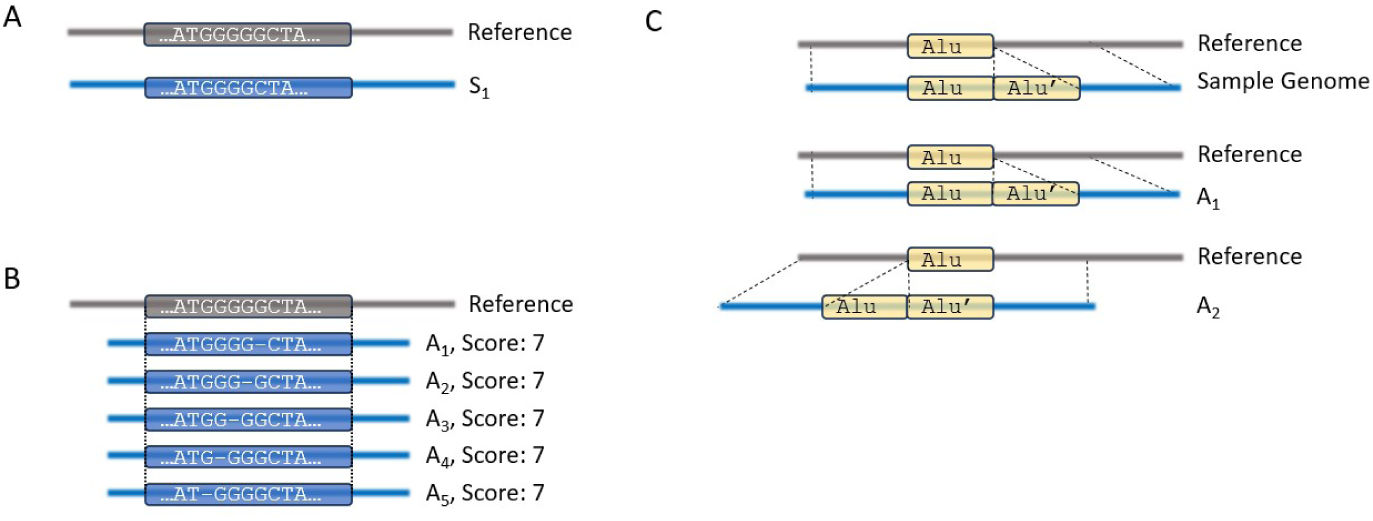
Alignment ambiguity may occur at different scales. **A)** A read, S1, and the reference where the length of the homopolymer run between the two read and the reference varies. **B)** Read S1 is aligned to the reference where the length of the homopolymer found in the read differs from the reference. There are multiple equivalent alignments of the read to the reference (*A*_1_ - *A*_5_) but with a varied deletion position. It is not clear which of the bases of the homopolymer run is deleted from the alignment of the read to the reference. All 5 alignments have the same alignment score. **C)** A new repeat copy occurs adjacent to an existing repeat element from the same repeat family in a sample. As there is very high sequence similarity between the two repeat copies, when aligning noisy long reads the aligner has to choose between two nearly equivalent alignments. This introduces ambiguity between the reads that support the repeat insertion as to where the exact location of the insertion copy is relative to the genome, upstream or downstream of the existing element.

To address both repeat alignment ambiguity and alignments truncated due to mobile element insertions we developed somrit, the somatic retrotransposon insertion toolkit, to detect novel somatic retrotransposon insertion events and MEIs from long reads mapped to a reference genome. Somrit is a modular toolkit consisting of subprograms with standard input/output files. Importantly, it has steps not found in traditional SV detection workflows aimed to recover insertions that may be missed due to alignment truncation, and to resolve repeat alignment ambiguity. In this work we first describe somrit, providing an overview of each sub-module and then show how somrit can be used to detect novel somatic MEIs and help avoid false positive translocations from general purpose SV callers.

## Methods

Somrit contains individual sub-modules designed to be run as standalone tools or as part of a larger workflow. **Figure 2** shows the somrit modules in the order they would normally be run to call somatic retrotransposon insertions.

**Figure 2.**
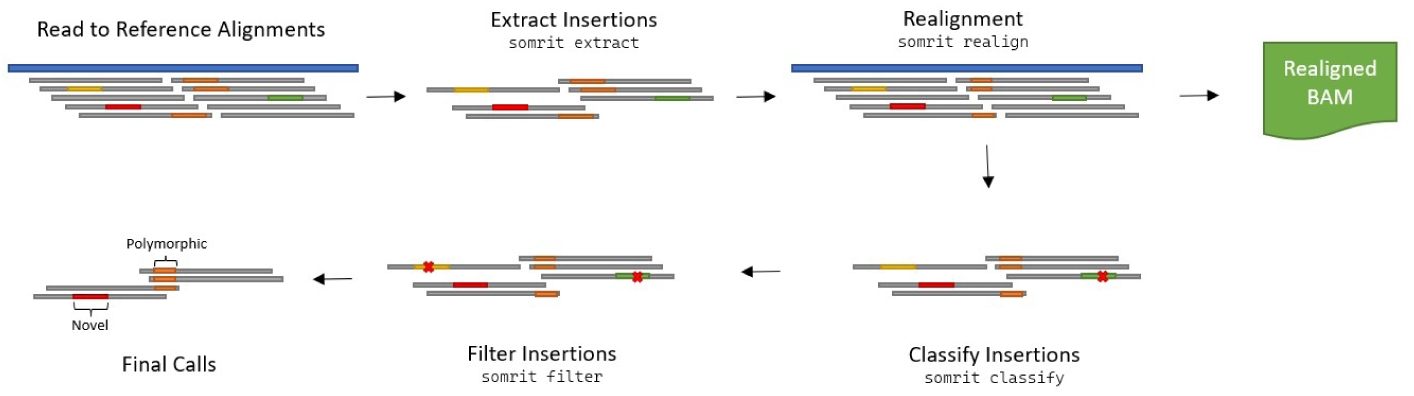
Somrit consists of sub-modules that can be run individually or in sequence. Somrit takes as input a BAM of reads aligned to a reference genome from a long read aligner such as minimap2. `somrit extract` identifies insertions in the long reads based on the alignments in the BAM. `somrit realign` performs local realignment to reduce genomic insert positional ambiguity and increase read support, outputting a realigned BAM. `somrit classify` annotates insertions as being from a retrotransposon repeat family while `somrit filter` applies a series of filters and annotations to identify insertions caused by mapping artifacts, those that may polymorphic, and those that have poly-A tails, target site duplications and LINE-1 endonuclease motifs.

`somrit extract.` Somrit’s first step is to extract candidate retrotransposon insertions from the reads aligned to the reference genome. We consider two cases. In the simple case, reads containing long insertions (by default, 50bp) with a minimum flanking anchor sequence (500bp) are exported to a tsv file. Second, we attempt to recover alignments that were erroneously split due to the presence of a large insertion within the read, shown in **Figure 3B**. Let *q.d, t.d* be the distance between the pair of alignments on the query (read) and target (reference), respectively. We merge the pair of alignments when *q.d ≥* 100 and *t.d ≤* 100 by writing the first BAM record with a new CIGAR string and deleting the second BAM record (**Figure 3C**). The coordinates of these insertions are also output to the TSV file.

**Figure 3.**
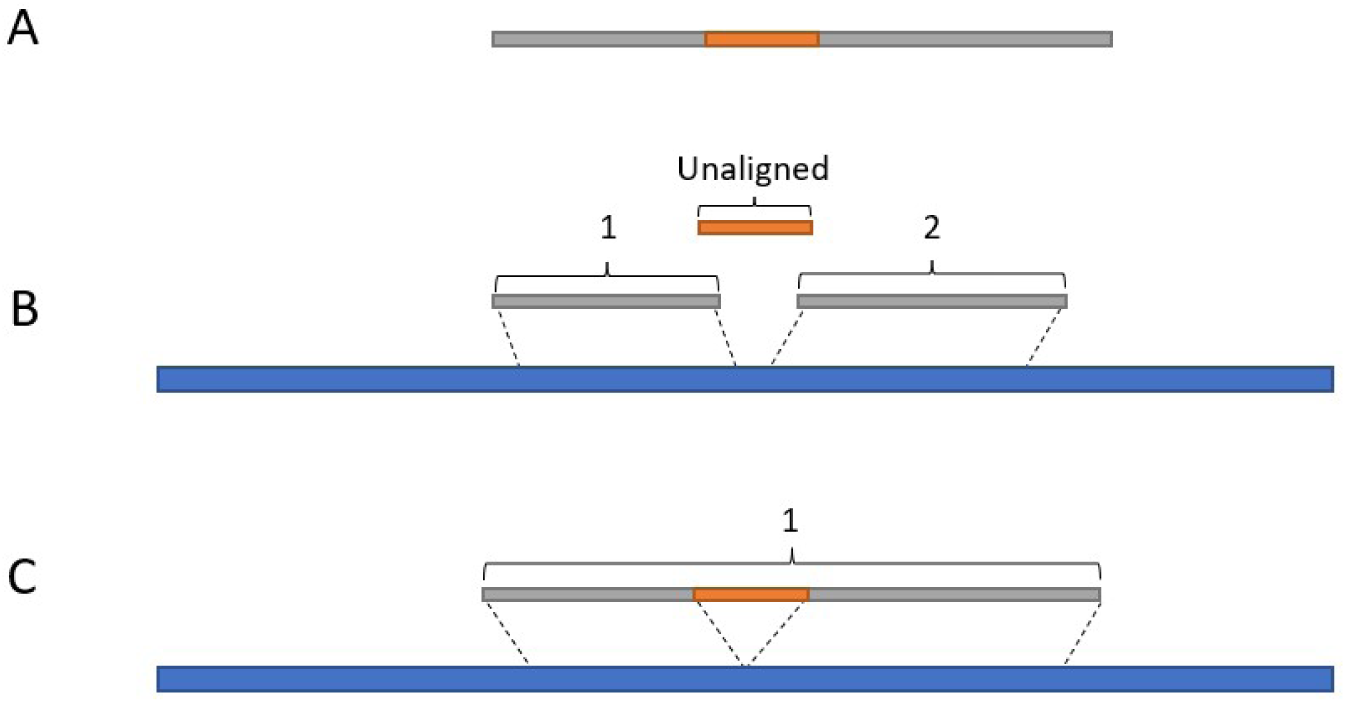
Merging of Split Alignments. **A)** A read containing a large insertion, shown in orange. **B)** The read is aligned to the reference (blue) with the sequence flanking the orange insertion aligned as two separate alignment records adjacent on the reference but with a larger gap on the read containing unaligned insertion sequence. **C)** After merging there is a single aligned segment for the read containing an insertion.

`somrit realign.` Next, we perform local realignment around candidate insertions to reduce alignment ambiguity and increase read support. The explicit goal of somrit is to detect novel repetitive insertions that have high sequence similarity to existing repeat copies within the reference genome. This can make it difficult for the read mapper to identify the correct insertion location when the insert happens to occur in a region already containing a copy of the repeat. In this case the output of `somrit extract` may have different representations of the same insertion event across multiple reads. As the level of read support is a key parameter for structural variant calling this can cause false negatives, or worse, the caller might identify multiple separate insertions. `somrit realign` aims to reconcile the alignments of all reads carrying an insertion and recover supporting reads entirely missed by the mapping and extract steps. This process is inspired by the predominant approach for small variant calling, which generates candidate haplotypes containing combinations of variants [41], [42], [43]. Here, we apply the same idea to large insertions found from long reads. `somrit realign` focuses on insertions at least *n* (default *n* = 50*bp*) employing a process similar to that of Iris [44], a tool for refining the position of structural variants in long reads, and SVJedi-Graph [45].

The `realign` module contains two steps: realignment and alignment projection. In the realignment step insertions identified by `somrit extract` (**Figure 4A**) are grouped based on genomic position into 1000bp windows (**Figure 4B**). Adjacent windows that contain insertions are merged together up to a max window size (default 25000bp). A set of consensus sequences (default=3, one for each germline haplotype for assumed diploid samples, and one for a germline haplotype containing the putative somatic insertion) is generated for each window from the insertion-supporting reads using abPOA [46]. Each consensus sequence is aligned back to the reference sequence for this window to identify a refined insertion position and sequence (**Figure 4C**). For each refined insert identified (minimize size 50bp), the insert is spliced into the reference to generate an alternative haplotype sequence that only contains a single insertion. Each read in the window is then aligned to the original reference as well as each alternative haplotype for the window. If the read’s alignment score against the alternative haplotype is greater than the alignment score to the reference we note the read as supporting the insertion and flag it for projection (**Figure 4D**).

**Figure 4.**
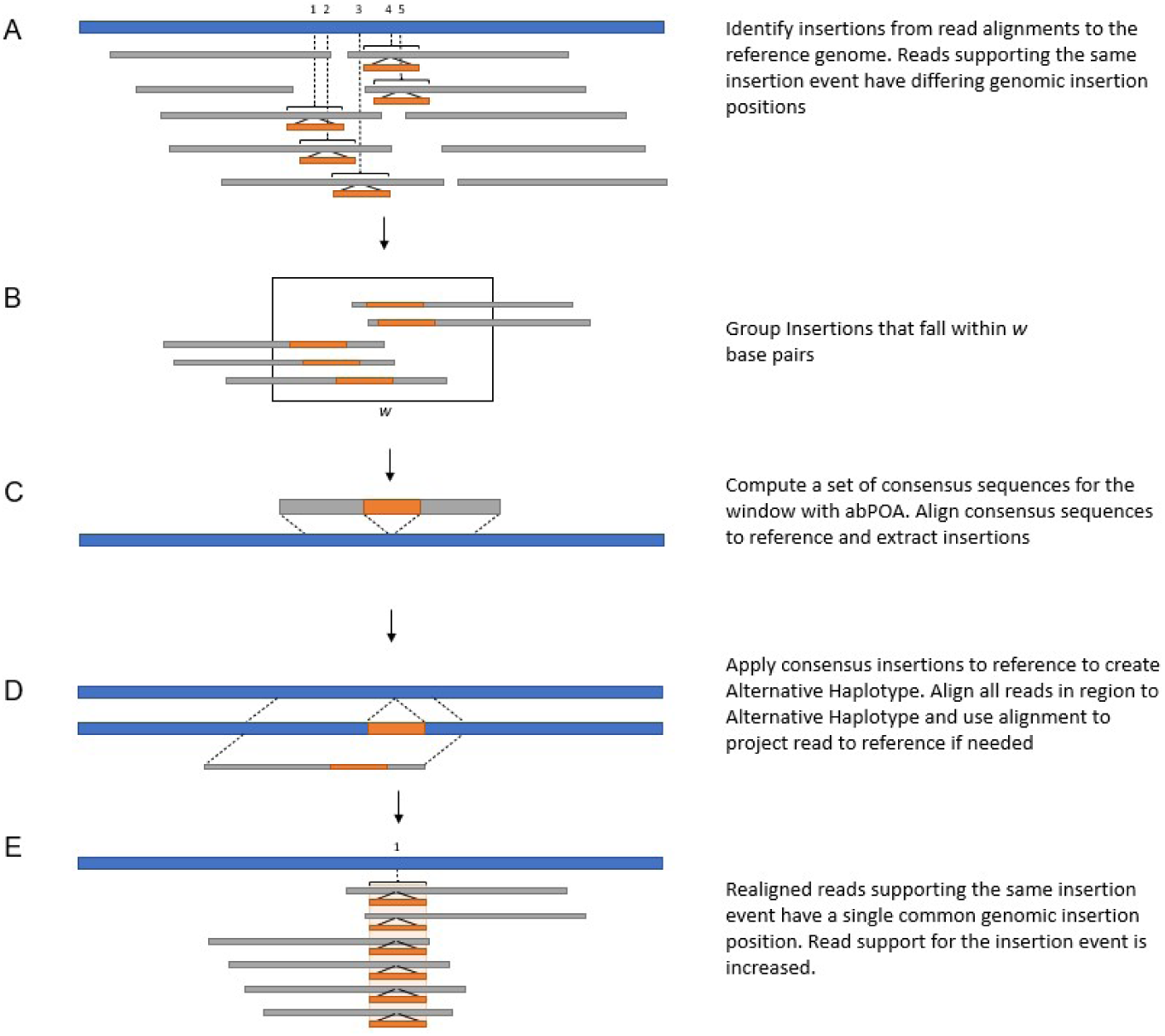
Somrit’s Realignment Process. **A)** Insertions supporting a single insertion event identified in the alignment of reads (grey) to the reference genome (blue) with `somrit extract` are shown in orange. These insertions have varying genomic insert positions, shown by the numbers above the blue reference track. It is not clear from the alignments of the reads where the exact insertion position is. **B)** Insertions are grouped together if they fall within *w* base pairs of each other on the reference. **C)** A set of consensus sequences is computed for the window, one for each haplotype present, with abPOA. These sequences are aligned to the reference and any insertions in the consensus sequences relative to the reference identified and extracted. **D)** Consensus insertions are applied to the reference genome to generate a set of alternative haplotypes containing insertion sequences. Reads are aligned to the alternative haplotypes to determine if they support the insertion contained within it. If deemed to support an insertion the alternative haplotype is used as a guide to project the read to the reference, allowing for more accurate placement of the insertion relative to the reference **E)** After projection and realignment the reads supporting the same insertion event now have a consistent genomic insertion position, with there being increased read support for the insertion event as well.

Once all reads have been tagged with the insertions they support, we calculate a new read-to-reference alignment in a step we call alignment projection. For each read a new haplotype is constructed by splicing in all insertions supported by that read. The read is then aligned to this haplotype, and the haplotype is aligned to the reference genome. We then iterate over the pair of read-to-haplotype and haplotype-to-reference CIGAR strings to determine the read-to-reference alignment. The BAM record for the read is then updated based on this projected alignment (**Figure 4E**). If a read is not selected for projection the original BAM record(s) for the read are retained. In addition to an updated BAM, `somrit realign` outputs an updated tsv with the coordinates and sequences of insertions after realignment.

`somrit classify.` The set of refined insertions are then assigned to a retrotransposon repeat family. Each insert’s sequence is aligned to a library of known human retrotransposon consensus sequences compiled from Tubio et al [27] (available: https://gitlab.com/mobilegenomesgroup/TraFiC) and DFAM [47] using minimap2’s mappy API. Inserts that have no mapping to a retrotransposon consensus sequence with quality higher than 20 are unassigned, otherwise the insert is assigned to the repeat family with the highest alignment score.

`somrit filter.` The final step applies annotations and filters to the classified repeats by appending new columns to the TSV record similar to VCF filter columns. These filters include:

- IN CONTROL SAMPLE: If somrit is run with multiple samples and one is designated a matched normal control sample (e.g. for tumour/normal pairs), this filter is used to identify which insertions are also found in the designated control sample within +/-500bp.
- IN CENTROMERE and IN TELOMORE: insertions that fall within a centromeric or telomeric region respectively based on a bed file provided by the user.
- LOW MAPPING QUALITY: Insertions found in a genomic window (+/- 500bp of the insertion position) where the average mapping quality over all reads aligned in the region is *≤* 20.
- MIN READS: This filter flags insertions that do not have the user-specified number of supporting reads (by default, 1).
- IN SECONDARY MAPPING: If reads supporting the insertion have multiple alignments with a mapping quality *≥* 20 that overlap the insertion position, the insertion is flagged.
- POLYMORPHIC: This filter flags insertions that appear to be polymorphic germline variation between the individual and the reference rather than somatic variation, based on the fraction of insertion supporting reads relative to all reads aligned in a genomic window. Let *f* (*w*) be the fraction of insertion supporting reads within a window of *w* bp. An insertion is flagged as polymorphic if *f* (500) *>* 0.8 or *f* (200) *>* 0.5 or *f* (100) *>* 0.3. The varied window sizes and fraction cutoffs were used as some genomic regions varied in coverage. A larger window may contain a number of reads that align within the window but do not overlap the insertion position, with parts of the window flanking the insertion having higher coverage than the area around the insertion itself.

In addition to these filters, each putative insertion is annotated with features expected of real retrotransposon insertions:

- Annotate Poly-A Tail: the insertion contains a Poly-A/T tail *≥* 10bp within 50bp of either the start or end of the insert sequence. If present the Poly-A/T sequence is listed in the output column.
- Annotate TSD: retrotransposon insertions have characteristic sequence duplication generated as part of the re-insertion process. A local dynamic programming alignment is used to identify duplicated sequence at least 5bp in length between the start or end of the insertion and the genomic region. If a duplication of at least 5bp is found, the TSD sequence is listed in the output column.
- Annotate Motif: the reference sequence 2bp upstream and 4bp downstream of the identified insertion position is extracted for comparison to the canonical LINE-1 endonuclease recognition motif sequence. The motif sequence is listed in the output column.

### Implementation and Pipeline

The tsv file generated by `somrit filter` is the final output of the program, with the inserts passing all filters considered the final called somatic insertions. Somrit is implemented python (`extract`, `classify` and `filter` modules) and C++ (`realign`). The code for all modules, a documentation of parameters, a tutorial, and a snakemake file to automate the process of running all 4 module sequentially are available at https://github.com/adcosta17/somrit.

## Results

In this section we first quantify how often repeat alignment ambiguity occurs. Next we evaluate somrit’s ability to detect both polymorphic and somatic insertions from simulated and real nanopore data, comparing somrit to existing tools for both tasks and showing how somrit’s use of local realignment to reduce repeat positional ambiguity and increase read support improved its ability to detect MEIs compared to other tools. Finally, we finally show how realignment around MEIs can reduce the number of false positive translocation calls from general purpose structural variation detection tools.

### 0.1 Repeat Alignment Ambiguity

We first quantified how often repeat alignment ambiguity occurs, specifically when a novel mobile element insertion occurs adjacent to an existing copy of the same mobile element in the genome. Using the RepeatMasker [48] annotation of GRCh38 we identified existing Alu elements at least 250bp in length. We then generated an insertion by randomly selecting an Alu element present in GRCh38, modifying it to a set level of sequence divergence and then inserting it back into the genome beside the original copy. This process simulates the insertion of an identical or near identical mobile element directly next to an existing mobile element. We then simulated long reads using pbsim2 [49], mean read length of 30kb and per-base accuracy of 95%, that supported the insertion and aligned the reads back to GRCh38 using minimap2 [36]. We parsed these read alignments to identify where the aligner had placed the insertion and compared it to the expected position where we had made the insertion. We generated 100 insertions for sequence divergence from 0-50%, in steps of 1%. If a read was mapped *>* 50bp away from the expected insertion position it was considered to be misaligned. We see from the results shown in **Figure 5** that at low levels of sequence divergence there is a high fraction of reads that misaligned, exhibiting repeat alignment ambiguity, as the aligner cannot differentiate between the existing copy and the new insertion copy. As sequence divergence increases the proportion of reads whose insertion is incorrectly placed decreases as the aligner is better able to differentiate between the repeat copies.

**Figure 5.**
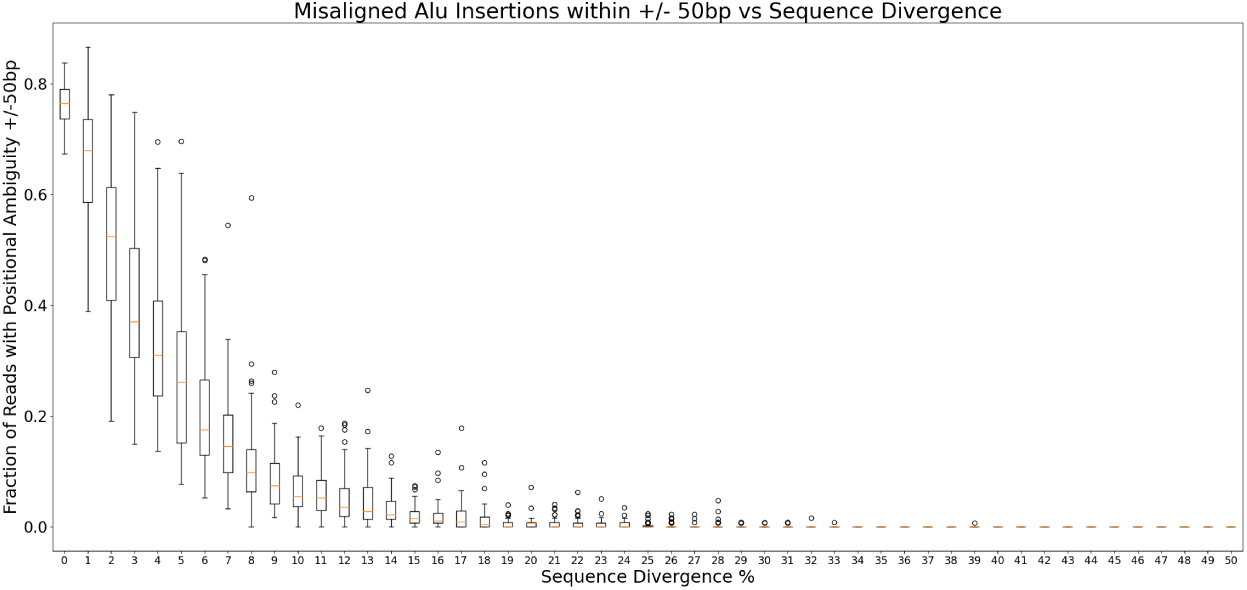
A mobile element insertion that occurs adjacent to an existing mobile element with high sequence similarity may result in a misaligned read. Using simulated reads that spanned insertions that are similar to an existing mobile element in the genome with a known sequence divergence, we measured the fraction of reads that contained an Alu insertion adjacent to an existing Alu element where the detected insertion position was off by *>* 50bp from the expected insertion position. We note that at low sequence divergence a higher fraction of reads are misaligned. As sequence divergence increased the fraction of misaligned reads decreased as the aligner is better able to differentiate between the repeat copies.

### Detection of Polymorphic Insertions

Polymorphic insertions, defined here as variants between an individual’s inherited genome and the reference, make up the vast majority of large insertions (*≥* 50bp) found by SV callers. To evaluate the performance of somrit in detecting polymorphic retrotransposon insertions compared to existing tools, xTea-Long and tldr, we used publicly available data downloaded from the Human Pan-genome Reference Consortium (HPRC)[35]. For samples HG00438, HG00621, HG00673, HG00735 and HG00741 we downloaded raw Oxford Nanopore (ONT) reads, the accompanying diploid hifiasm assembly [50] and matching RepeatMasker [48] annotation of the assembled contigs. The ONT reads for each of the five samples were downsampled to set coverage levels, with three replicates drawn for each coverage level. These read sets were then aligned to GRCh38 and the resulting BAM files passed as input to somrit, xTea-Long and tldr.

#### Generating ground truth calls

We used the high quality diploid assemblies for each HPRC sample to derive a set of ground truth insertions. For each sample the maternal and paternal contigs were aligned to GRCh38 with minimap2 (v2.22-r1101, preset `asm5`). Insertions at least 100bp in length on contig alignments with mapping quality *≥* 20 were identified and extracted from the alignment CIGAR strings. These insertions were then annotated with repeat families using the RepeatMasker [48] annotation of the contigs. Insertions where at least half the insertion sequence on the contig was annotated to a single retrotransposon repeat family by RepeatMasker were assigned to that repeat family. These insertions, their repeat family annotation (if any), and their reference coordinate are our truth set for the subsequent evaluation.

#### Comparing somrit, tldr and xTea-Long

We ran somrit with default settings except for increasing the minimum insertion supporting read threshold to 3 and requiring at least 1000bp of flanking sequence on each side of an insertion on the read. Also, we did not use somrit’s polymorphic filtering step. We ran tldr (v1.2.2) and xTea-Long (v0.19) using their default parameters, except for also requiring at least 1000bp of flanking sequence for tldr. We compared the retrotransposon insertion calls made by each tool to the ground truth described in the previous section. A called insertion was considered a true positive if it is within 500bp of an insertion call from the same retrotransposon repeat family in the truth set. All other called insertions were considered false positives. Any insertions in the truth set where we did not find a called insertion within 500bp annotated to the same retrotransposon repeat family were considered false negatives.

**Figure 6** shows the precision and recall for all three tools at different coverage levels, with 3 replicates per sample per coverage level. Somrit has higher precision at lower coverage and slightly lower precision at higher coverage compared to xTea-Long and tldr. At higher levels of coverage false insertions from mapping artifacts may have their read support increased beyond the threshold of 3 supporting reads, resulting in false positive calls. xTea-Long had higher recall than somrit and tldr at the lowest coverage levels, while tldr and somrit had higher recall than xTea-Long at higher coverage levels, with somrit’s recall slightly higher than tldr at the highest coverage levels. While tldr considers both reads that fully span an insertion event and reads whose alignments are clipped at the insertion event as supporting the insertion, somrit only looks at reads with a spanning insertion. This may contribute to the observed recall for the somrit being lower than tldr at lower coverage levels.

**Figure 6.**
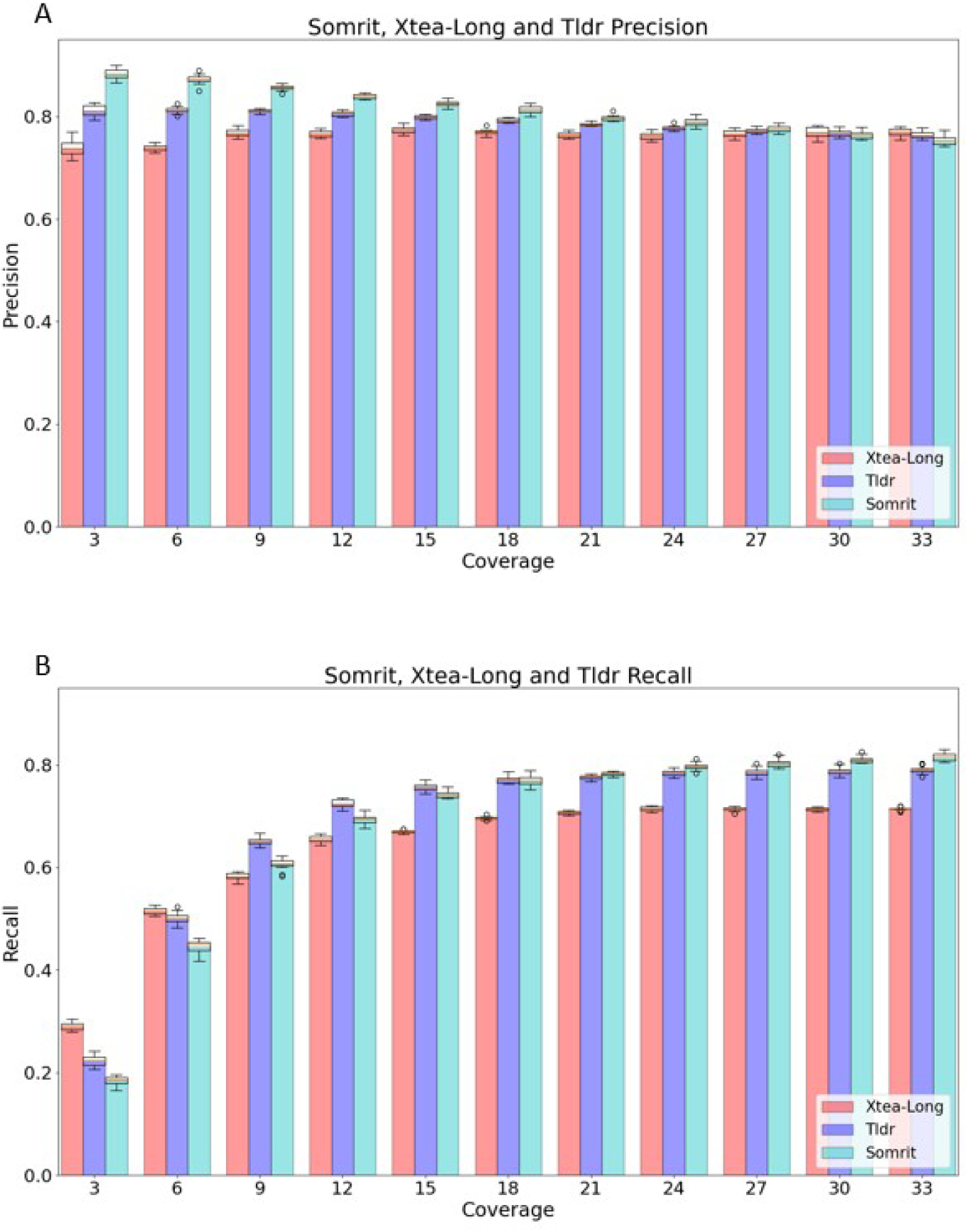
Somrit, XTea-Long and Tldr Precision (panel A) and Recall (panel B) for the detection of polymorphic retrotransposon insertions over 5 HPRC samples at varying coverage levels

### Detection of simulated somatic insertions

While somrit can be used to detect polymorphic insertions it is designed primarily to detect somatic insertions. This is more challenging than detecting polymorphic insertions as the read support may be much lower, even down to a single read. While tools like tldr have been previously used to detect rare somatic retrotransposon insertions[51], they are not designed for detecting insertions from a single read, which is one design goal of somrit. To quantify somrit’s ability to detect novel somatic retrotransposon insertion events we ran somrit on a set of simulated novel somatic retrotransposon insertions. For comparison we ran tldr and xTea-Long with the minimum read support lowered to 1. We also ran the general purpose SV caller Sniffles2 (v2.0) [52] in its somatic detection mode.

#### Generating Simulation Data

We simulated long reads with pbsim2[49] from the diploid assembly for 4 of the 5 HPRC samples: HG00438, HG00621, HG00735 and HG00741. We simulated 20x coverage from both the maternal and paternal contigs (40x total), to act as a baseline read set free from somatic variation (**Figure 7A** and **B**). Next, we randomly selected 500 positions on the assembly contigs for the location of somatic insertions. For each selected position we randomly choose a retrotransposon repeat family (LINE-1, Alu or SVA) and insert length. For half of the selected positions a full length insertion is selected with the length of the remainder drawn uniformly between 100 bp and the full repeat length. Using a consensus sequence for the repeat family selected [27][47], we truncated the sequence if needed removing bases from the 5’ end, generated a poly-A tail at the 3’ end between 10 and 40 bp and added the modified sequence to the contig at the selected position. We also generated a target site duplication (TSD) to mimic the real genomic insertion process. We then simulated long reads from each contig in a 50kbp flanking region around the insertion position. We recorded which reads fully spanned the insertion with non-insert flanking sequence. For each position on the contig where we added an insertion we noted the insert sequence, repeat family, poly-A tail length, TSD, and the expected location of the insertion on GRCh38, and the list of read names deemed to support the insertion event (**Figure 7C**). In order to generate each test read set that contained simulated somatic retrotransposon insertion events we started with the simulated baseline read set for the sample and randomly selected 125 of the 500 positions where we had simulated a novel somatic insertion event. For each of these 125 positions we randomly selected between 1 to 4 supporting reads and added them to the base read set to generate a test read set (**Figure 7D** and **E**). This process is repeated to generate 12 replicates per sample, each with a randomly generated set of 125 positions.

**Figure 7.**
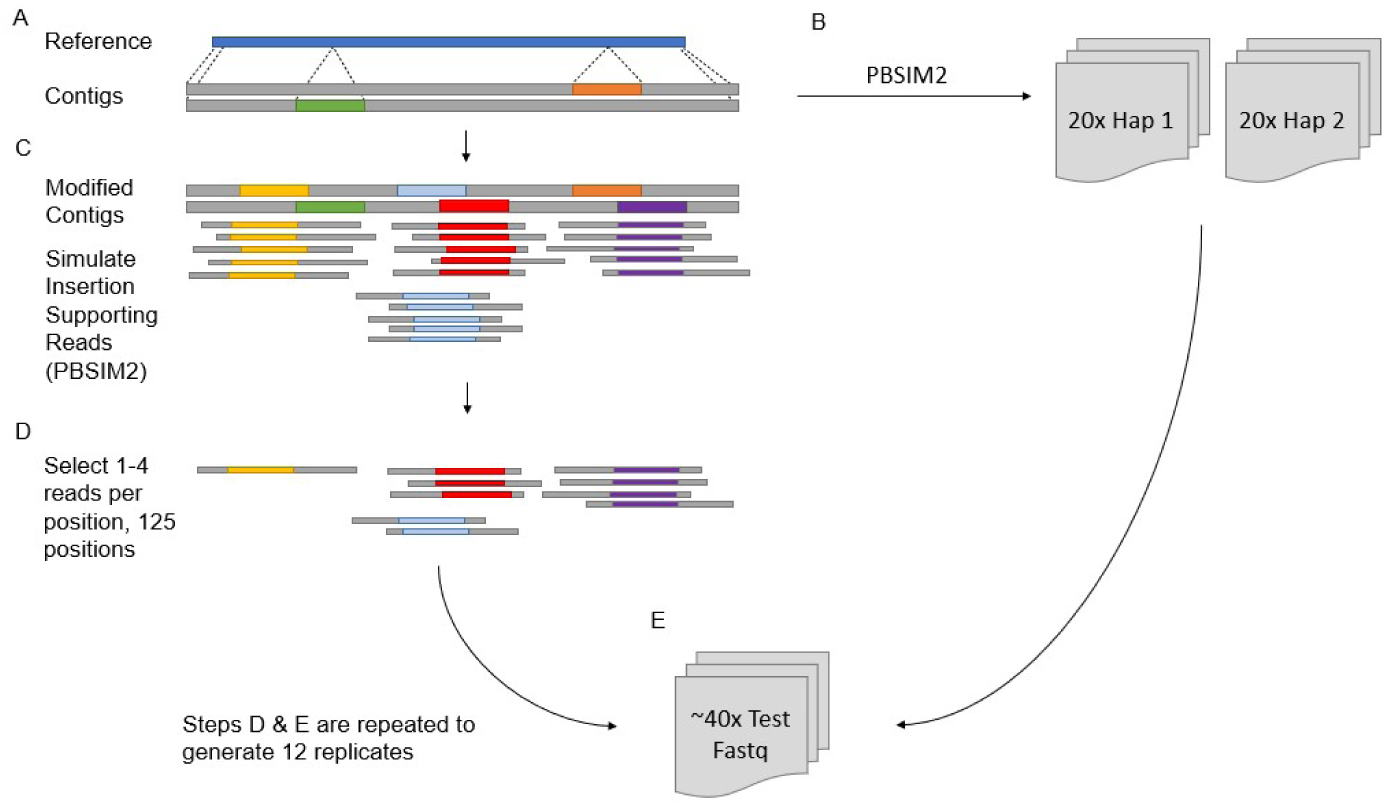
Generating Simulated Somatic Insertions. **A)** The diploid assembly of an HPRC sample is aligned to the reference to identify where each contig aligns to. **B)** pbsim2 is used to generate two 20x base read sets of simulated ONT reads, one for haplotype in the diploid assembly. These read sets represent normal reads with only germline polymorphic variation. **C)** Insertions are added to the contigs at 500 randomly selected positions to mimic novel retrotransposon insertion events. Insertions are randomly selected to be from either LINE-1, Alu or SVA repeat families and to be partial or full length with a poly-A tail and target site duplication included. Reads are simulated at these regions with reads that span the added insertion identified **D)** 125 positions where insertions have been made are randomly chosen, and 1 to 4 reads per position that support the insertion selected. **E)** These insertions are added to the two base read sets generated in step B to generate an 40x test read set. Steps D and E are repeated 12 times to generate 12 replicates.

#### Detecting simulated insertion events

For each replicate we mapped the reads to GRCh38 with minimap2 and then ran tldr, Sniffles2 and somrit as described above to identify somatic insertions in the same 4 HPRC samples. We ran tldr and somrit in their default settings but with the minimum read flank size set to 1000bp and minimum read support set to 1. We compared the calls made by each tool to the truth data for each replicate, noting the insertion events that were detected as passing retrotransposon insertions by somrit and tldr as well any insertions detected by Sniffles2 (as Sniffles2 does not annotate insertions as being from a retrotransposon repeat family). Passing insertion calls made within 500bp of an expected simulated insertion position with the same repeat family annotation were considered true positives. If no passing insertion with the same repeat annotation was found within 500bp of a simulated insertion position, the simulated insertion position was considered a false negative. We additionally ran xTea-Long in its default settings but as xTea-Long is not designed for the detection of somatic insertions it was unable to detect almost all the simulated insertions. Thus we do not report the results of xTea-Long in this analysis.

We computed recall for each tool and over all replicates for the 4 HPRC samples (**Figure 8**). Somrit had the highest recall, followed by Sniffles2. As somrit generates calls for individual reads so we also calculate a read-level recall (the proportion of reads, rather than positions, that have an insertion that were called by somrit; **Figure 8 inset**). Even though somrit outperformed all other tools, its best recall for any sample did not exceeded 60%, indicating the difficulty of detecting insertions with minimal read support. Due to the repetitive nature of the human genome a repeat insertion event in a long read makes it harder for a long read aligner to correctly align the read. Thus reads with insertions may either have split alignments at the expected insertion position where realignment is unable to increase the read support, or may not align to the expected region of the reference, or the reference at all. All tools evaluated rely on alignment BAM files as input and thus are limited by the shortcomings of current long read aligners when aligning reads with repeat insertions.

**Figure 8.**
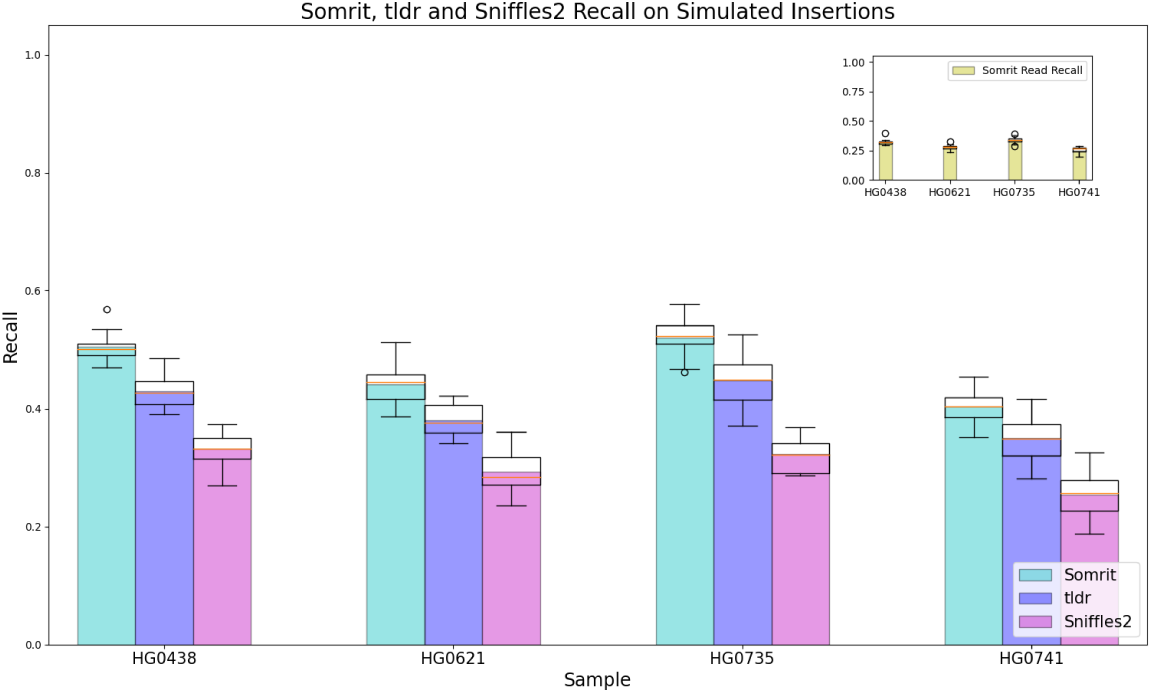
Somrit, Tldr and Sniffles2 Simulated Insertion Recall. The recall of somrit, tldr and Sniffles2 for detecting simulated somatic retrotransposon insertions over 4 HPRC samples, 12 replicates per sample. Each simulated somatic insertion event is supported by 1 to 4 reads, ranges in length from 100 to 6000 base pairs and includes LINE-1, Alu or SVA sequence. As somrit provides a file containing read support for each insertion call we also computed recall at the level of individual sequencing reads, shown in the top right inset plot.

### Identifying novel L1-mCherry retrotransposon insertions in Nanopore reads

In a recent analysis by Gerdes et al [51] HeLa cells were treated with a plasmid vector containing a modified mouse LINE-1 fused with an mCherry reporter. Successful integration of this modified vector into the HeLa cells results in a novel insertion of the L1-mCherry construct sequence in the cells. This is an ideal experiment to test the performance of somatic retrotransposon insertion detection tools as the mCherry sequence allows novel insertions to be definitively identified. Using the ONT data generated by Gerdes et al for these samples we evaluated somrit’s ability to call these previously identified insertions.

We downloaded reads from the HeLa/L1-mCherry experiment that Gerdes et al deposited in the ENA. The construct generated insertions of up to 10.2kbp once integrated into the HeLa genome[51]. Each of the five read sets are WGS nanopore sequencing run of a HeLa cell line expanded over 3-5 passages from single L1-mCherry insertion harboring colonies, barcoded and pooled in equal amounts before being sequenced with a single PromethION flow cell [51]. It is expected that individual L1-mCherry insertions will appear as somatic insertion events occurring in a small fraction of cells in the sample, with possibly just a single read supporting the insertion event.

We ran somrit and tldr on the 5 samples, with minimum read support set to 1 and a minimum read flank size of 1000 bp. We ran both tools using the L1-mCherry consensus as the only possible repeat family. Both tools were run in multi-sample mode, where multiple samples are analysed jointly. This allowed for insertion calls to have supporting reads from multiple biological replicate cell line colonies, making it easier to identify novel somatic events that only occur in a single sample rather than those that represent any possible polymorphic variation seen across all samples. As the L1-mCherry sequence contained both a full length LINE-1 sequence and the mCherry protein coding gene we explicitly filtered the final detected insertion calls to ensure they aligned with at least one base pair to the non LINE-1 sequence of the L1-mCherry construct.

The results in **Table 1** show that somrit is able to identify additional insertions containing L1-mCherry sequence beyond what was initially detected by Gerdes et al. Each of the insertion calls made by somrit used *≤* 5 reads, with 37 insertions being detected with just a single supporting read. Both tools detected insertions that were unique to the tool. The 5 insertions unique to tldr represent the 5 of the 6 insertions of the Gerdes et al set that somrit did not identify. Of these 5 somrit was able to identify 3 as being annotated to the L1-mCherry sequence, but these insertions were flagged for not having enough flanking sequence in the read alignment. Manual inspection of the 10 insertions unique to somrit showed that 8 of the 10 were mapped to the 3’ end of the L1-mCherry construct sequence, with target site duplications and poly-A tails or mapped to non-LINE1 sequence in the L1-mCherry construct, indicating they are likely to be novel insertions of the L1-mCherry construct sequence into the cells. The remaining two insertions called only by somrit mapped mainly to the LINE1 portion of the L1-mCherry construct, with only a small fraction of the insertion sequence aligning to non-LINE-1 sequence in the construct, with no mCherry specific sequence identified. Thus these insertions are likely false positives.

**Table 1.** L1-mCherry insertions detected by Somrit and Tldr. The total number of L1-mCherry insertions, the number of insertions unique to each tool, the number of insertions shared between the tools and the number of insertions shared with Gerdes et al’s call set.

|  | Total Inserts | Unique To Tool | Shared Between Tools | Shared with Gerdes et al |
| --- | --- | --- | --- | --- |
| somrit | 67 | 10 | 57 | 35/41 |
| tldr | 62 | 5 | 57 | 40/41 |

### Repeat realignment reduces false positive translocation calls

While somrit is primarily designed to detect mobile element insertions, local realignment with `somrit realign` may be useful in reducing false positive calls from general purpose SV callers such as Sniffles2 and CuteSV. As the human genome contains many copies of mobile repetitive elements such as retrotransposons and the exact location of these elements varies between individuals and the reference genome, some sequencing reads that partially cover a non-reference mobile repetitive element may appear to have a split mapping when aligned to the reference genome. This split mapping occurs as the aligner may map the non-repetitive sequence correctly but maps the portion of the read containing the mobile element sequence to an existing repetitive element copy elsewhere in the genome. If a number of reads are misaligned in this way a general purpose SV caller may incorrectly interpret this as a translocation. We propose that `somrit realign` may help reduce the number of false positive translocation calls induced by this effect (**Figure 9**).

**Figure 9.**
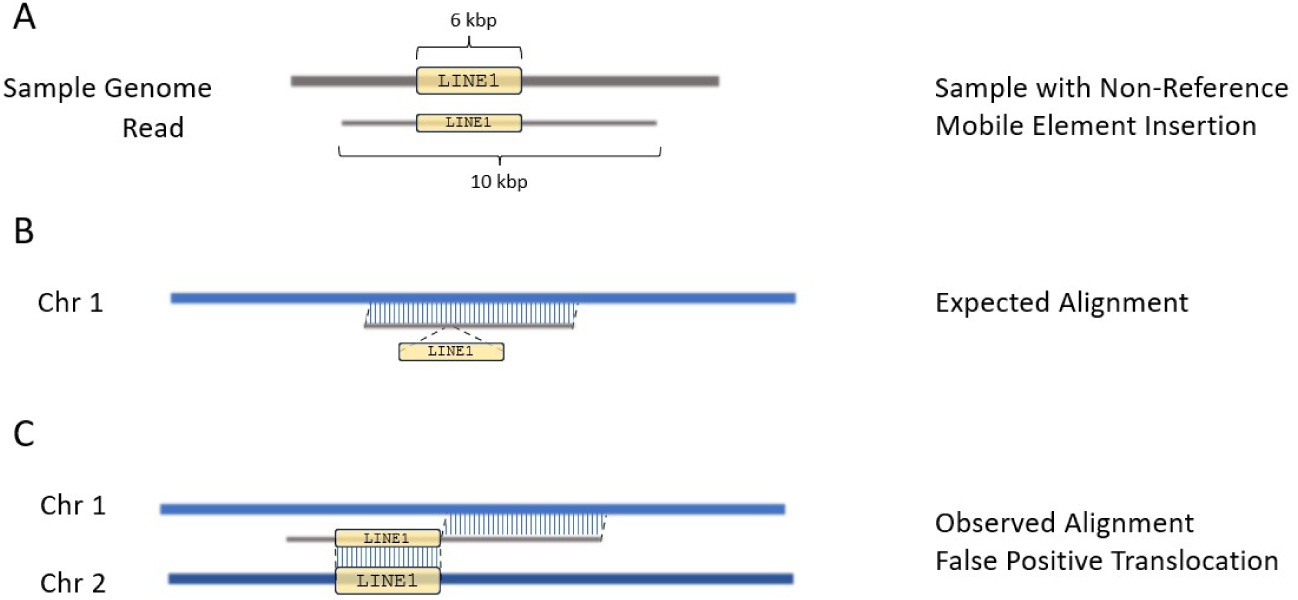
Non-Reference Mobile Elements can cause false positive translocation calls A) A read originating from a sample that contains a non-reference LINE1 insertion and a read that fully spans the insertion. B) When this read is aligned to the reference genome we expect an alignment containing an insertion gap relative to the chromosome 1 reference sequence. C) Alternatively, the read may be aligned as two distinct segments. In the first segment, the non-repeat sequence matches chromosome 1 at the expected position. In the second segment, the LINE1 mobile element insertion sequence aligns to an existing LINE1 copy elsewhere in the genome on a different chromosome. If enough reads share the same pattern of split mapping a false positive translocation may be called between these two positions.

To evaluate how `somrit realign` could be used to reduce false positive translocation calls we first ran Sniffles2 and CuteSV on the aligned reads for each HPRC sample at various read depths, noting the number of inter-chromosomal translocation calls made. As the HPRC samples are generated from lymphoblastoid cell lines classified as karyotypically normal, thus free of any known inter-chromosomal translocation events, we considered any inter-chromosomal translocation calls made by the tools as false positives. We then detected candidate insertions (`somrit extract`) for realignment (`somrit realign`) to generate a new BAM file. We then used the realigned bam as input into Sniffles2 and CuteSV, noting the number of called inter-chromosomal translocations after realignment.

**Figure 10** shows that in both tools there is a reduction in the number of false positive inter-chromosomal translocation calls made after realignment. We observe up to 41% and 31% reduction in the number of false positive inter-chromosomal translocation calls made by Sniffles2 and CuteSV, respectively, with the effect most noticeable at higher coverage levels.

**Figure 10.**
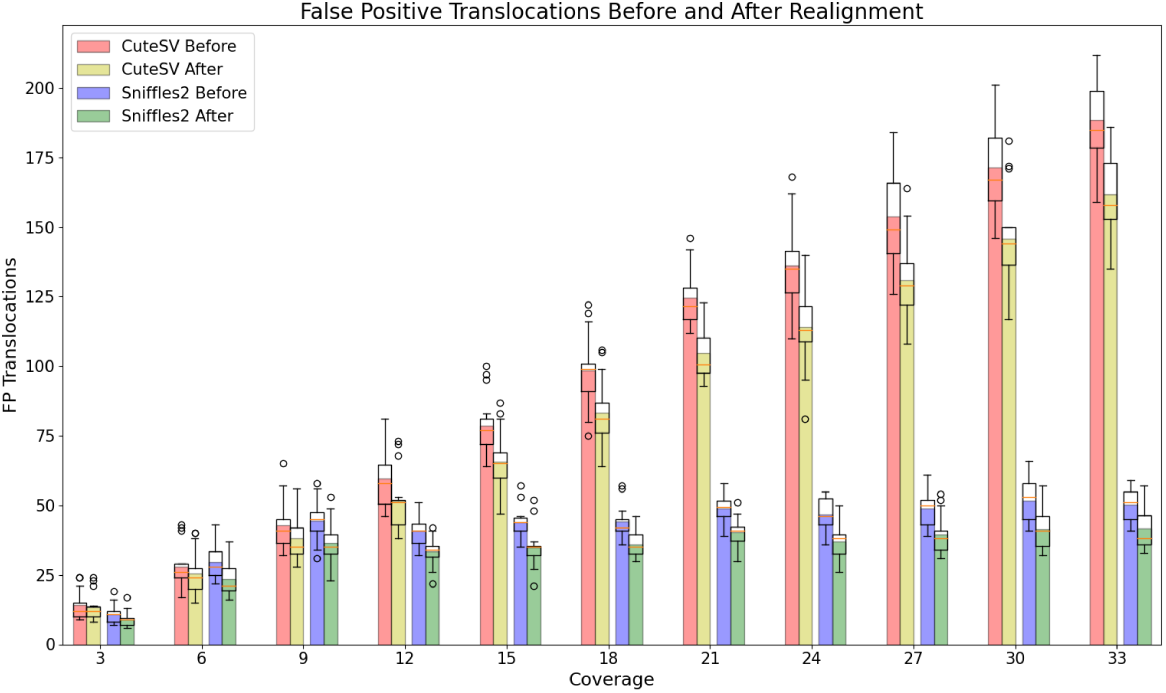
Sniffles and CuteSV False Positive Inter-Chromosomal Translocation calls. The number of false positive inter-chromosomal translocation calls made by Sniffles and CuteSV before and after Realignment. Each down-sampled fastq from the 5 normal HPRC samples assumed to be free of any known translocations relative to GRCh38 was passed to Sniffles and CuteSV before and after realignment, with the number of called translocations made by each tool shown.

## Discussion

In this paper we introduce somrit, a toolkit for the identification of somatic retrotransposon insertion events in long reads. We show that somrit is able to detect existing polymorphic MEIs with comparable precision and recall to state-of-the-art tools. We also show that somrit is able to detect somatic MEIs in both simulated and real nanopore data, outperforming other methods at identifying insertions with single read support. In addition, we show that realignment around MEIs can reduce false positive translocation calls in general purpose SV callers.

While these results show somrit’s effectiveness, they have limitations. Somrit firstly requires a large amount of time and memory to run, more than existing tools for retrotransposon insertion detection. The majority of the computational burden lies with `somrit realign` and the generation of consensus sequences with abPOA. While abPOA uses adaptive banded alignment to reduce the time and memory usage needed to compute a consensus sequence this process is still time and memory intensive. As the time and memory needed to compute the consensus sequences scales with both the number and length of input sequences, limiting the number of read sequences used to generate the consensus sequences at high input coverage can be considered to reduce the time and memory usage.

Somrit is also limited in its ability to realign insertions that may be missed by the initial mapping of reads to the reference. If there is an insertion present in a sample, but there is no alignment made by the aligner that introduces an alignment gap for the insertion, somrit is unable to recover this insertion. This becomes problematic for somatic detection where insertions may have low read support and an aligner may clip the alignment of the single read supporting an insertion event, with somrit unable to detect or recover the insertion.

While realignment is able to increase the read support for genuine insertion events, in some cases the realignment process may increase the number of reads that support a mapping or alignment artifact. The decision to realign a read using an alternative haplotype containing an insertion as a guide is based on comparing the alignment score between the alternative and reference haplotypes. A higher scoring alignment to the alternative haplotype indicates that the read may support the insertion. If an false alternative haplotype is generated by a mapping artifact, and the read has a marginally higher alignment score to this haplotype, a mapping artifact could be introduced into the read, with the read now supporting a false insertion. We believe this effect can be seen in the decreased precision somrit has compared to other tools at higher levels of coverage for polymorphic insertion detection, as at higher coverage levels there is a greater chance a mapping artifact supported by one or two reads has its read support increased through realignment to three or more reads. More stringent criteria for selecting alternative haplotypes may help alleviate this issue.

## Acknowledgements

We thank the members of the Simpson Lab at the Ontario Institute for Cancer Research for their helpful suggestions, ideas and feedback in debugging. A.D. was the recipient of an Ontario Graduate Scholarship. J.T.S receives funding from the Government of Ontario through the Ontario Institute of Cancer Research.

## Code Availability

Somrit is available at https://github.com/adcosta17/somrit. Scripts to generate the evaluation and analysis performed as well as generate the plots shown in this paper can be found at https://github.com/adcosta17/somrit-test.

## Supplementary Information

### Time and Memory Analysis

We evaluated how somrit compared to other SV detection methods for computational performance. We noted the total time and memory usage of somrit, xTea-Long, tldr and Sniffles2 runs during the previously mentioned analysis of simulated somatic retrotransposon insertion events on the 4 HPRC samples. The analysis of simulated somatic retrotransposon insertion events used a number of different machines as part of a larger shared computing environment, thus tools were not run on the same machine. While not ideal this approach does allow us to get a range of possible time and memory usages for a tool over different machines with equivalently sized input. For each HPRC sample each tool was run the 12 40x replicates used for the simulated insertion analysis.

Figure 11 shows this comparison for memory usage and Figure 12 for time. As somrit consists of multiple individual modules, we noted the time of each step and reported two versions of the time analysis. One version, shown in Figure 12 as somrit total, represents the total time taken if each step is run sequentially with 10 threads. The second version, shown in Figure 12 as somrit ideal, is the total time taken if individual re-alignment jobs for each chromosome are run in parallel with 10 threads each. For somrit memory we took the maximum memory over all modules for a given input fastq.

**Figure 11.**
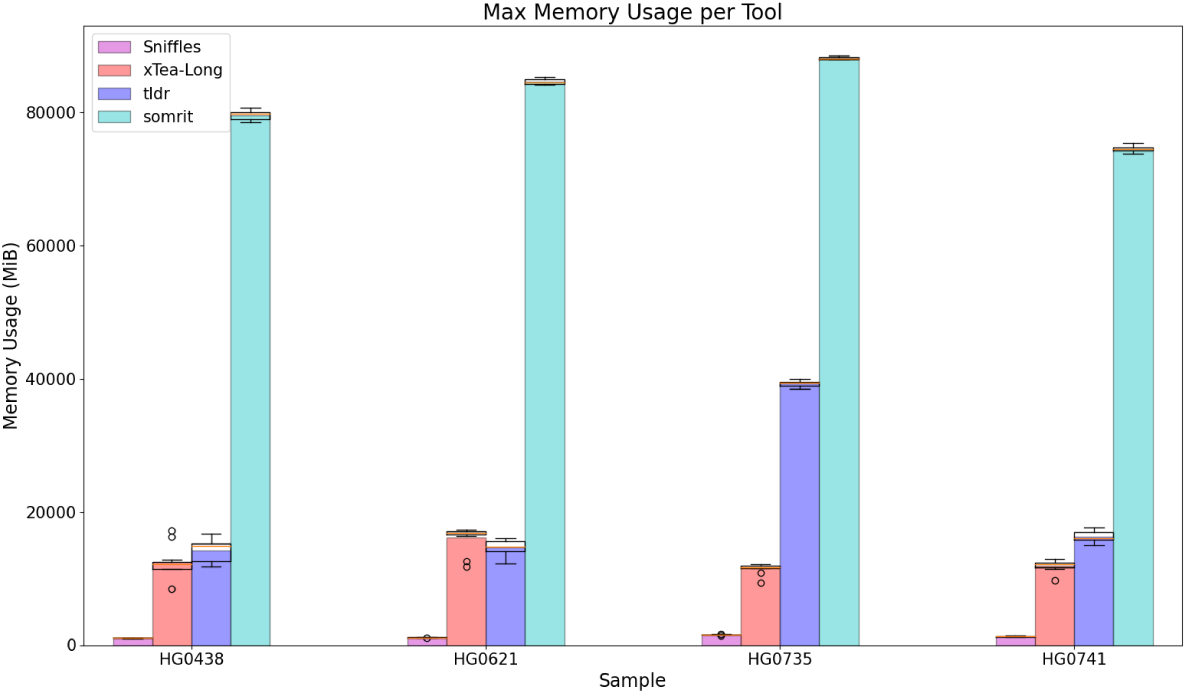
Memory Usage per Tool. The maximum memory usage of somrit, xTea-Long, tldr and Sniffles2 for each replicate and coverage level over five HPRC samples.

**Figure 12.**
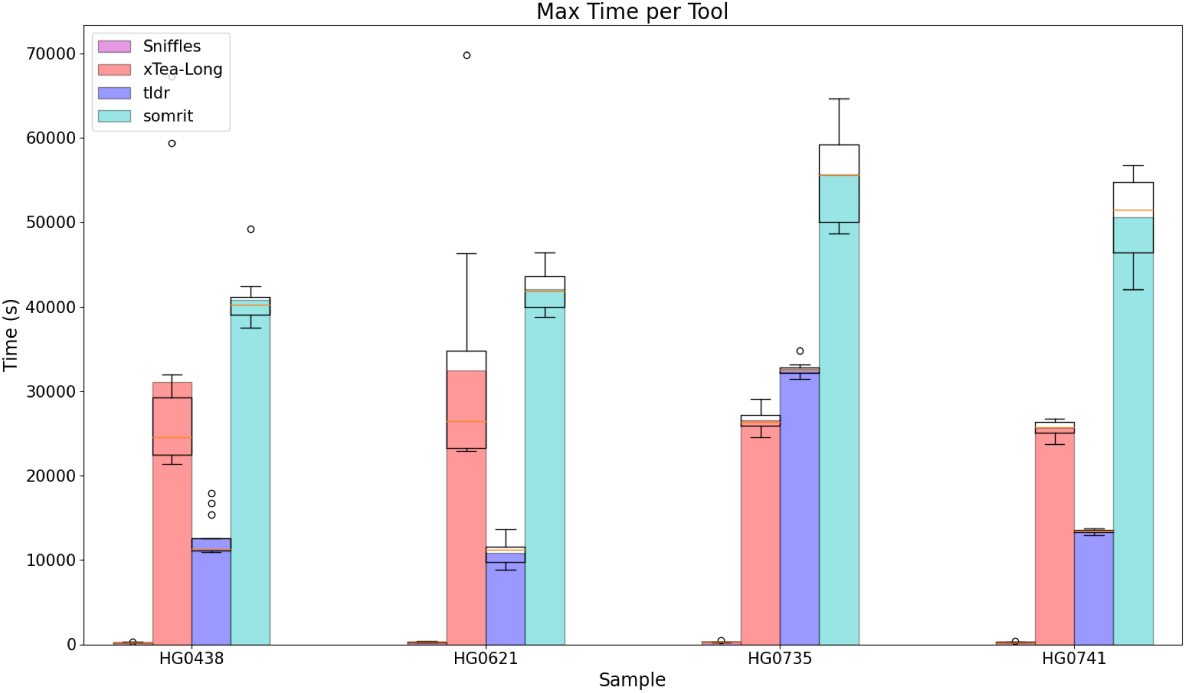
Elapsed Time per Tool. The elapsed time for somrit, xTea-Long, tldr and Sniffles2 run on each replicate and coverage level over five HPRC samples.

Somrit does have both higher run time overall and higher memory usage than other tools at 40x coverage. If somrit realign is run in parallel per chromosome the total time required for somrit is comparable to that of tldr and xTea-Long. The higher memory usage of somrit is attributed to the consensus generation step of realignment, with abPOA requiring a high amount of memory to generate consensus sequences.

## References

1. Bourque, G., Burns, K.H., Gehring, M., Gorbunova, V., Seluanov, A., Hammell, M., Imbeault, M., Izsvák, Z., Levin, H.L., Macfarlan, T.S., et al.: Ten things you should know about transposable elements. Genome biology 19(1), 1–12 (2018))

2. Ostertag, E.M., Kazazian Jr, H.H.: Biology of mamalian l1 retrotransposons. Annual review of genetics 35, 501 (2001)

3. Szak, S.T., Pickeral, O.K., Makalowski, W., Boguski, M.S., Landsman, D., Boeke, J.D.: Molecular archeology of l1 insertions in the human genome. Genome biology 3(10), 1–18 (2002))

4. Sudmant, P.H., Rausch, T., Gardner, E.J., Handsaker, R.E., Abyzov, A., Huddleston, J., Zhang, Y., Ye, K., Jun, G., Hsi-Yang Fritz, M., et al.: An integrated map of structural variation in 2,504 human genomes. Nature 526(7571), 75–81 (2015))

5. Penzkofer, T., Dandekar, T., Zemojtel, T.: L1base: from functional annotation to prediction of active line-1 elements. Nucleic acids research 33(suppl 1), 498–500 (2005))

6. Beck, C.R., Collier, P., Macfarlane, C., Malig, M., Kidd, J.M., Eichler, E.E., Badge, R.M., Moran, J.V.: Line-1 retrotransposition activity in human genomes. Cell 141(7), 1159–1170 (2010))

7. Hancks, D.C., Kazazian, H.H.: Roles for retrotransposon insertions in human disease. Mobile DNA 7(1), 1–28 (2016))

8. Hancks, D.C., Kazazian Jr, H.H.: Active human retrotransposons: variation and disease. Current opinion in genetics & development 22(3), 191–203 (2012))

9. Goodier, J.L.: Restricting retrotransposons: a review. Mobile DNA 7(1), 1–30 (2016))

10. Xiao-Jie, L., Hui-Ying, X., Qi, X., Jiang, X., Shi-Jie, M.: Line-1 in cancer: multifaceted functions and potential clinical implications. Genetics in Medicine 18(5), 431–439 (2016))

11. Dombroski, B.A., Mathias, S.L., Nanthakumar, E., Scott, A.F., Kazazian Jr, H.H.: Isolation of an active human transposable element. Science 254(5039), 1805–1808 (1991))

12. Dewannieux, M., Esnault, C., Heidmann, T.: Line-mediated retrotransposition of marked alu sequences. Nature genetics 35(1), 41–48 (2003))

13. Ade, C., Roy-Engel, A.M., Deininger, P.L.: Alu elements: an intrinsic source of human genome instability. Current opinion in virology 3(6), 639–645 (2013))

14. Ostertag, E.M., Goodier, J.L., Zhang, Y., Kazazian Jr, H.H.: Sva elements are nonautonomous retrotransposons that cause disease in humans. The American Journal of Human Genetics 73(6), 1444–1451 (2003))

15. Feng, Q., Moran, J.V., Kazazian Jr, H.H., Boeke, J.D.: Human l1 retrotransposon encodes a conserved endonuclease required for retrotransposition. Cell 87(5), 905–916 (1996))

16. Symer, D.E., Connelly, C., Szak, S.T., Caputo, E.M., Cost, G.J., Parmigiani, G., Boeke, J.D.: Human l1 retrotransposition is associated with genetic instability in vivo. Cell 110(3), 327–338 (2002))

17. Moran, J.V., Holmes, S.E., Naas, T.P., DeBerardinis, R.J., Boeke, J.D., Kazazian Jr, H.H.: High frequency retrotransposition in cultured mammalian cells. Cell 87(5), 917–927 (1996))

18. Doucet, A.J., Wilusz, J.E., Miyoshi, T., Liu, Y., Moran, J.V.: A 3 poly (a) tract is required for line-1 retrotransposition. Molecular cell 60(5), 728–741 (2015))

19. Pickeral, O.K., Makałowski, W., Boguski, M.S., Boeke, J.D.: Frequent human genomic dna transduction driven by line-1 retrotransposition. Genome research 10(4), 411–415 (2000))

20. Goodier, J.L., Ostertag, E.M., Kazazian Jr, H.H.: Transduction of 3-flanking sequences is common in l1 retrotransposition. Human molecular genetics 9(4), 653–657 (2000))

21. Ewing, A.D., Kazazian, H.H.: High-throughput sequencing reveals extensive variation in human-specific l1 content in individual human genomes. Genome research 20(9), 1262–1270 (2010))

22. Cajuso, T., Sulo, P., Tanskanen, T., Katainen, R., Taira, A., Hänninen, U.A., Kondelin, J., Forsström, L., Välimäki, N., Aavikko, M., et al.: Retrotransposon insertions can initiate colorectal cancer and are associated with poor survival. Nature communications 10(1), 1–9 (2019))

23. Han, K., Lee, J., Meyer, T.J., Remedios, P., Goodwin, L., Batzer, M.A.: L1 recombination-associated deletions generate human genomic variation. Proceedings of the national academy of sciences 105(49), 19366–19371 (2008))

24. Sen, S.K., Han, K., Wang, J., Lee, J., Wang, H., Callinan, P.A., Dyer, M., Cordaux, R., Liang, P., Batzer, M.A.: Human genomic deletions mediated by recombination between alu elements. The American Journal of Human Genetics 79(1), 41–53 (2006))

25. Scott, E.C., Gardner, E.J., Masood, A., Chuang, N.T., Vertino, P.M., Devine, S.E.: A hot l1 retrotransposon evades somatic repression and initiates human colorectal cancer. Genome research 26(6), 745–755 (2016))

26. Gardner, E.J., Lam, V.K., Harris, D.N., Chuang, N.T., Scott, E.C., Pittard, W.S., Mills, R.E., Devine, S.E., Consortium, .G.P., et al.: The mobile element locator tool (melt): population-scale mobile element discovery and biology. Genome research 27(11), 1916–1929 (2017))

27. Tubio, J.M., Li, Y., Ju, Y.S., Martincorena, I., Cooke, S.L., Tojo, M., Gundem, G., Pipinikas, C.P., Zamora, J., Raine, K., et al.: Extensive transduction of nonrepetitive dna mediated by l1 retrotransposition in cancer genomes. Science 345(6196), 1251343 (2014))

28. Keane, T.M., Wong, K., Adams, D.J.: Retroseq: transposable element discovery from next-generation sequencing data. Bioinformatics 29(3), 389–390 (2013))

29. Chu, C., Borges-Monroy, R., Viswanadham, V.V., Lee, S., Li, H., Lee, E.A., Park, P.J.: Comprehensive identification of transposable element insertions using multiple sequencing technologies. Nature communications 12(1), 1–12 (2021))

30. Marsili, L., Duque, K.R., Bode, R.L., Kauffman, M.A., Espay, A.J.: Uncovering essential tremor genetics: The promise of long-read sequencing. Frontiers in neurology 13 (2022)

31. Pollard, M.O., Gurdasani, D., Mentzer, A.J., Porter, T., Sandhu, M.S.: Long reads: their purpose and place. Human molecular genetics 27(R2), 234–241 (2018))

32. Gong, T., Hayes, V.M., Chan, E.K.: Detection of somatic structural variants from short-read next-generation sequencing data. Briefings in Bioinformatics 22(3), 056 (2021))

33. Jain, M., Olsen, H.E., Paten, B., Akeson, M.: The oxford nanopore minion: delivery of nanopore sequencing to the genomics community. Genome biology 17(1), 1–11 (2016))

34. Ewing, A.D., Smits, N., Sanchez-Luque, F.J., Faivre, J., Brennan, P.M., Richardson, S.R., Cheetham, S.W., Faulkner, G.J.: Nanopore sequencing enables comprehensive transposable element epigenomic profiling. Molecular Cell 80(5), 915–928 (2020))

35. Wang, T., Antonacci-Fulton, L., Howe, K., Lawson, H.A., Lucas, J.K., Phillippy, A.M., Popejoy, A.B., Asri, M., Carson, C., Chaisson, M.J., et al.: The human pangenome project: a global resource to map genomic diversity. Nature 604(7906), 437–446 (2022))

36. Li, H.: Minimap2: pairwise alignment for nucleotide sequences. Bioinformatics 34(18), 3094–3100 (2018))

37. Jain, C., Rhie, A., Hansen, N.F., Koren, S., Phillippy, A.M.: Long-read mapping to repetitive reference sequences using winnowmap2. Nature Methods 19(6), 705–710 (2022))

38. Jain, C., Rhie, A., Zhang, H., Chu, C., Walenz, B.P., Koren, S., Phillippy, A.M.: Weighted minimizer sampling improves long read mapping. Bioinformatics 36(Supplement 1), 111–118 (2020))

39. Ren, J., Chaisson, M.J.: lra: A long read aligner for sequences and contigs. PLOS Computational Biology 17(6), 1009078 (2021))

40. Audano, P.A., Beck, C.R.: Small allelic variants are a source of ancestral bias in structural variant breakpoint placement. bioRxiv, 2023–06 (2023))

41. Albers, C.A., Lunter, G., MacArthur, D.G., McVean, G., Ouwehand, W.H., Durbin, R.: Dindel: accurate indel calls from short-read data. Genome research 21(6), 961–973 (2011))

42. Poplin, R., Ruano-Rubio, V., DePristo, M.A., Fennell, T.J., Carneiro, M.O., Van der Auwera, G.A., Kling, D.E., Gauthier, L.D., Levy-Moonshine, A., Roazen, D., et al.: Scaling accurate genetic variant discovery to tens of thousands of samples. BioRxiv, 201178 (2018)

43. Garrison, E., Marth, G.: Haplotype-based variant detection from short-read sequencing. arXiv preprint arXiv:1207.3907 (2012))

44. Kirsche, M., Prabhu, G., Sherman, R., Ni, B., Aganezov, S., Schatz, M.C.: Jasmine: Population-scale structural variant comparison and analysis. BioRxiv, 2021–05 (2021))

45. Romain, S., Lemaitre, C.: Svjedi-graph: improving the genotyping of close and overlapping structural variants with long reads using a variation graph. Bioinformatics 39(Supplement 1), 270–278 (2023))

46. Gao, Y., Liu, Y., Ma, Y., Liu, B., Wang, Y., Xing, Y.: abpoa: an simd-based c library for fast partial order alignment using adaptive band. Bioinformatics 37(15), 2209–2211 (2021))

47. Storer, J., Hubley, R., Rosen, J., Wheeler, T.J., Smit, A.F.: The dfam community resource of transposable element families, sequence models, and genome annotations. Mobile DNA 12(1), 1–14 (2021))

48. Chen, N.: Using repeat masker to identify repetitive elements in genomic sequences. Current protocols in bioinformatics 5(1), 4–10 (2004))

49. Ono, Y., Asai, K., Hamada, M.: Pbsim2: a simulator for long-read sequencers with a novel generative model of quality scores. Bioinformatics 37(5), 589–595 (2021))

50. Cheng, H., Concepcion, G.T., Feng, X., Zhang, H., Li, H.: Haplotype-resolved de novo assembly using phased assembly graphs with hifiasm. Nature methods 18(2), 170–175 (2021))

51. Gerdes, P., Lim, S.M., Ewing, A.D., Larcombe, M.R., Chan, D., Sanchez-Luque, F.J., Walker, L., Carleton, A.L., James, C., Knaupp, A.S., et al.: Retrotransposon instability dominates the acquired mutation landscape of mouse induced pluripotent stem cells. Nature Communications 13(1), 1–18 (2022))

52. Smolka, M., Paulin, L.F., Grochowski, C.M., Mahmoud, M., Behera, S., Gandhi, M., Hong, K., Pehlivan, D., Scholz, S.W., Carvalho, C.M., et al.: Comprehensive structural variant detection: from mosaic to population-level. Biorxiv, 2022–04 (2022))

